# Genomic characterization of six virus-associated cancers identifies changes in the tumor microenvironment and altered genetic programming

**DOI:** 10.1101/289900

**Authors:** Frederick S. Varn, Yue Wang, Chao Cheng

## Abstract

Viruses affect approximately 20% of all human cancers and induce the expression of viral oncoproteins that make these tumors potent targets for immune checkpoint inhibitors. In this study, we apply computational tools to The Cancer Genome Atlas and other datasets to define how virus infection shapes the tumor microenvironment and genetic architecture of 6 virus-associated tumor types. Across cancers, the cellular composition of the microenvironment varied by viral status, with infected tumors often exhibiting increased infiltration of cytolytic cell types. Analyses of the infiltrating T cell receptor repertoire revealed that Epstein-Barr virus was associated with decreased diversity in multiple cancers, suggesting an antigen-driven immune response. Tissue-specific gene expression signatures capturing these virus-induced transcriptomic changes successfully predicted virus status in independent datasets and were associated with both immune- and proliferation-related features that were predictive of prognosis. The analyses presented suggest viruses have distinct effects in different tumors with implications for immunotherapy.

## Introduction

Immune checkpoint inhibitors have yielded promising results in treating cancer. By targeting the proteins that attenuate T cell receptor (TCR) signaling following antigen recognition, these drugs can initiate a robust anti-tumor immune response that induces remission and prolongs survival in subgroups of patients with multiple malignancies.^1-7^ Despite the early successes of these treatments, response rates to these therapies remain low, creating a need for biomarkers that are capable of identifying responsive patients. Tumor mutation burden has been identified as one such biomarker, with tumors exhibiting high mutation loads more likely to express immunogenic neoantigens that are recognized as non-self by the adaptive immune system.^8^ This association has been supported across tumor types, with high-mutation tumors yielding higher response rates than those tumors with lower mutation rates^9,10^ and also within tumor types, with responsive patients exhibiting significantly higher mutation burdens.^11-16^ Given these associations, it is possible that other sources of antigen in the tumor microenvironment function similarly with regards to responses to immunotherapy.

Virus infections are a source of tumor antigen that affect approximately 20% of all human cancers. There are seven virus families that have been implicated as oncogenic, including hepatitis virus B and C (HBV and HCV), human papillomavirus (HPV), human herpesvirus 4, also known as Epstein–Barr virus (HHV4), human T-cell lymphotropic virus (HTLV), Merkel cell polyomavirus (MCV), and human herpesvirus 8, also known as Kaposi’s sarcoma virus (HHV8).^17^ HPV, HTLV, HHV4, MCV, and HHV8 are known to directly contribute to oncogenesis by expressing oncogenic proteins encoded in their genome. Conversely, HBV and HCV are involved in indirect carcinogenesis, causing chronic inflammation of the infected organs.^18^ Like neoantigens, the proteins expressed by these viruses are recognized as foreign by the immune system, with infection associated with increased immune activity in some tumor types.^19,20^ Immune checkpoint inhibitors thus provide a potentially effective treatment option for patients with these virus-induced cancers. However, the role viral proteins play in the magnitude and quality of these responses, and how these roles differ across cancer types, remains unclear. Two clinical trials testing the anti-PD-1 agent pembrolizumab in head and neck cancer and Merkel cell carcinoma found higher response rates among virus-positive patients compared to their virus-negative counterparts.^21,22^ Moreover, in Merkel cell carcinoma, MCV-positive patients exhibited significantly lower mutation burdens than virus-negative ones, indicating that viral antigen may be a sufficient immunogen to induce response to anti-PD1.^22^ Similar results were shown in gastric cancer, where a study identified an HHV4-positive patient responsive to anti-PD-L1 despite having a low tumor mutation burden.^23^ However, a study in cervical cancer revealed that the antitumor T cell response in HPV-positive tumors receiving adoptive T cell therapy was directed against a patient’s cancer germline antigens or neoantigens rather than viral antigens.^24^

Several studies have now characterized the occurrence of virus infections in thousands of tumor samples from The Cancer Genome Atlas (TCGA).^25,26^ Additionally, recent advances in genome-based immune profiling technologies have made it possible to systematically characterize the tumor microenvironment of large cohorts of patients. Several methods exist that use gene expression information to infer the infiltration level of different immune cell types in the tumor microenvironment.^20,27,28^ Other methods have been designed that use sequencing data contained in raw RNAseq reads to profile the TCR and B cell receptor repertoires from bulk tumor datasets.^29,30^ These methods together enable large-scale analyses that are more highly-powered to detect associations than traditional studies using flow cytometry and targeted receptor sequencing-based approaches. This is especially beneficial when studying virus infection in different tumor types, as these infections are exceedingly rare in some contexts.

In this study, we apply these tools to TCGA and other datasets to define how virus infection shapes the immune response in six tumor types: bladder urothelial carcinoma (BLCA), cervical squamous cell carcinoma (CESC), colorectal adenocarcinoma (COADREAD), head and neck squamous cell carcinoma (HNSC), liver hepatocellular carcinoma (LIHC) and stomach adenocarcinoma and esophageal carcinoma (STES). We begin by examining how the infiltration levels of CD8+ T cells, B cells, natural killer (NK) cells, and macrophages vary based on the presence of a virus infection in each tumor. We then perform TCR sequencing on bulk tumor data to assess how TCR diversity changes in relation to infection with different viruses and determine the extent to which virus infections are associated with clonal T cell responses. To expand our study beyond datasets that include virus infection information, we develop a gene signature to predict virus infection status in each of the tumor types in our study. We then functionally characterize this signature and apply it in a survival-meta analysis to provide a multi-dataset consensus on how virus infection affects cancer-specific patient prognosis. Together, these analyses provide novel insights into the altered tumor-immune dynamics associated with virus infection that can be used to refine our understanding of virus-associated tumor immunogenicity.

## Results

### Virus infection is associated with an altered tumor microenvironment

To obtain a global overview of how the tumor microenvironment differs in relation to virus infection, we applied our previously developed computational framework^27^ to infer immune infiltration levels of four distinct immune cell types, CD8+ T cells, B cells, NK cells, and macrophages, from TCGA RNAseq gene expression profiles of six virus-associated tumor types (Supplementary Table S1). We then stratified patients from each tumor type based on whether they were positive for reads from at least one virus, as determined by a prior study (Figure 1A).^26^ In 4/6 tumor types, infection was associated with elevated levels of CD8+ T cell infiltration, with CESC and HNSC exhibiting significant differences (P = 0.01 and 4e-5, respectively). We observed similar trends for B cells, with significant differences in BLCA, CESC, and HNSC (P = 0.01, 4e-3, and 2e-8, respectively, two-tailed Wilcoxon sum-rank test), as well as for NK cells, with significant elevations in the virus-positive samples of CESC, HNSC, and STES (P = 5e-3, 0.02, and 3e-4, respectively). Conversely, we observed significantly reduced abundance in macrophages when comparing HNSC samples (P = 2e-5).

**Figure 1:**
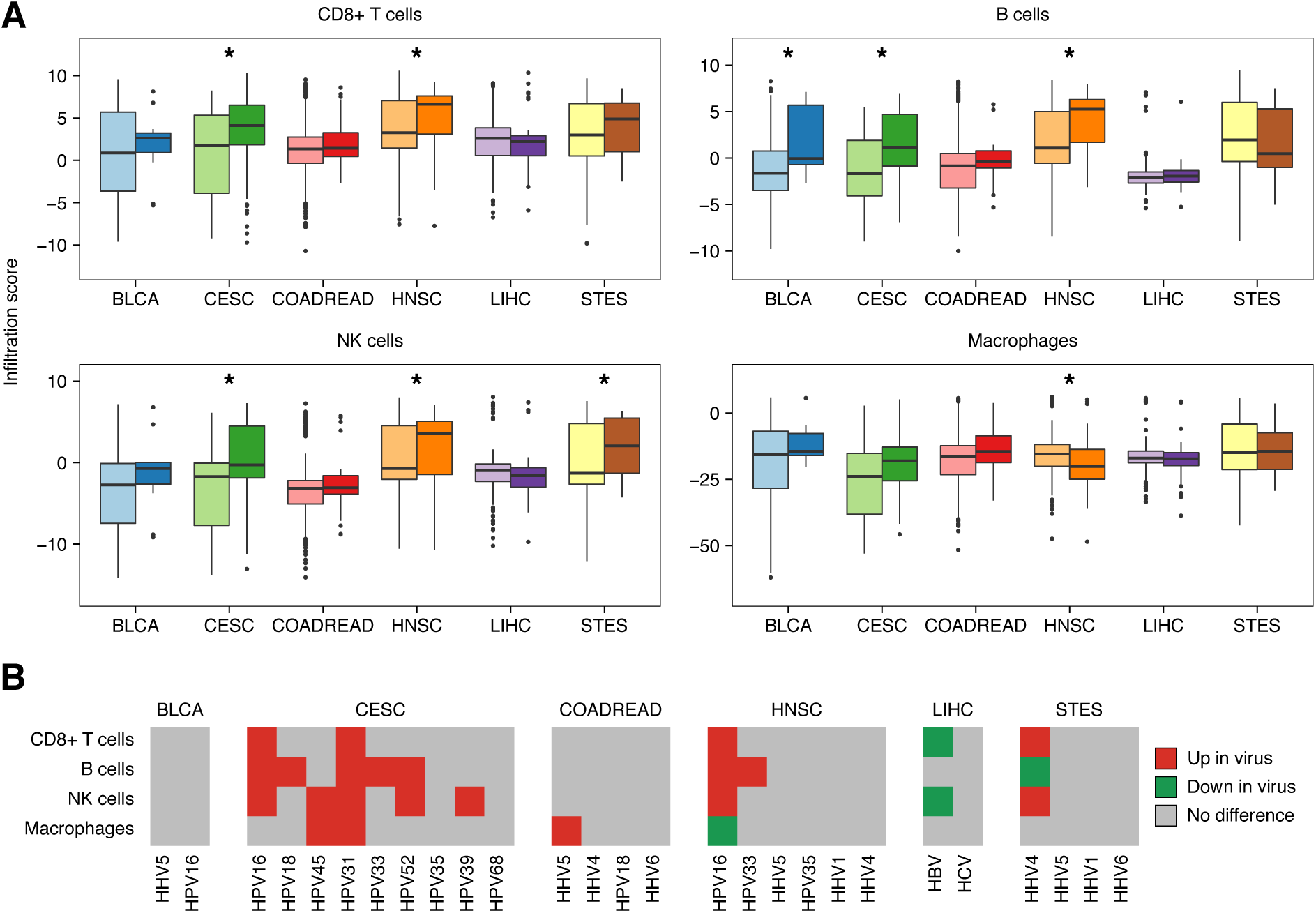
Differences in immune infiltration levels between virus infected and non-virus infected samples. **A**, Boxplots depicting the distribution of immune cell infiltration scores across samples from six cancer types stratified by virus infection status. Dark colors indicate virus-infected samples and light colors indicate non-infected samples. Each box spans quartiles with the lines representing the median infiltration score for each group. Whiskers represent absolute range excluding outliers. All outliers were included in the plot. Significant associations are marked (* p < 0.05). **B,** Heatmap marking significant differences in immune infiltration scores between samples infected with noted viruses and non-infected samples. All viruses infecting more than one patient in the denoted tumor type are shown. Red color indicates significant increases in infected samples (p < 0.05), green indicates significant decreases (p < 0.05) and grey indicates no significant difference (p > 0.05). All P-values were calculated using the two-tailed Wilcoxon sum-rank test.

We next examined how the type of virus affected theses associations. For this analysis, we further stratified patients based on the specific virus they were infected with and then compared each subgroup’s level of immune infiltrate to the virus-negative subgroup (Figure 1B). In several cases, there were only a small number of patients with a specific virus infection in a given tumor type, making it difficult to detect associations. However, in most tumor types, there were sufficient numbers of infections from at least one virus to detect significant differences in immune infiltrate. We found that in CESC, numerous viruses were associated with elevated infiltration levels of distinct cell types, with the most common virus, HPV16, associated with significantly higher levels of CD8+ T cells, B cells, and NK cells. In HNSC which also exhibits frequent HPV16 infection, we observed similar associations, including increased CD8+ T cell, B cell and NK cell infiltration as well as decreased macrophage infiltration. In STES, where samples were most frequently infected by HHV4, we observed significantly elevated CD8+ T cell and NK cell levels and decreased levels of B cell infiltration in HHV4-positive samples. Interestingly, these associations were reversed in HBV-infected LIHC, with HBV-positive samples exhibiting significantly lower levels of CD8+ T cell and NK cell infiltration. These samples also exhibited decreased expression of HLA-class I genes, suggesting the loss of antigen presentation machinery in these cancers (P = 0.07, 0.02, and 0.01 for HLA-A, HLA-B, and HLA-C, respectively; Supplementary Figure S1). These divergent results indicate that viruses of different families alter the tumor microenvironment of the samples they infect in distinct ways.

### Virus-associated immune infiltration associations are confounded by microsatellite instability

Microsatellite instability (MSI) is a condition associated with a high mutation burden that is a result of defects in the DNA mismatch repair pathway. This condition is especially prevalent in colorectal, gastric, and endometrial tumor types^31^ and has been shown to be associated with high levels of CD8+ T cell infiltration.^27^ We hypothesized that the observed differences in CD8+ T cell infiltration between virus infected and non-infected COADREAD and STES samples may have been confounded by the microsatellite instability of a patient’s tumor, masking potential increases in infiltration between virus-positive and virus-negative samples. To assess the extent to which this was true, we stratified patients from the COADREAD and STES cohorts into four groups depending on their MSI and virus infection status. We found that in both tumor types the majority of virus-positive samples were in the microsatellite stable (MSS) subgroup, with this subgroup containing 31/33 virus-positive samples in COADREAD and 50/55 samples in STES. Interestingly, in COADREAD the virus-positive/MSS samples exhibited significantly higher levels of CD8+ T cell infiltration compared to their virus-negative/MSS counterparts (P = 0.05, one-tailed Wilcoxon sum-rank test; Figure 2), while MSI samples exhibited significantly higher CD8+ T cell levels than either MSS group, regardless of virus status (P = 2e-14 and 9e-4, for virus-negative and virus-positive, respectively). However, in STES these associations were weaker, with virus-positive/MSS samples showing insignificant increases compared to virus-negative samples (P = 0.06, one-tailed Wilcoxon sum-rank test) and MSI samples exhibiting significantly higher levels of CD8+ T cell infiltration compared to the MSS/virus-negative samples, but not the MSS/virus-positive samples (P = 3e-3 and 0.34, respectively; Figure 2). Together, these results provided preliminary evidence that MSI status may confound associations between CD8+ T cell infiltration and other potential immunogenic factors in the microenvironment.

**Figure 2:**
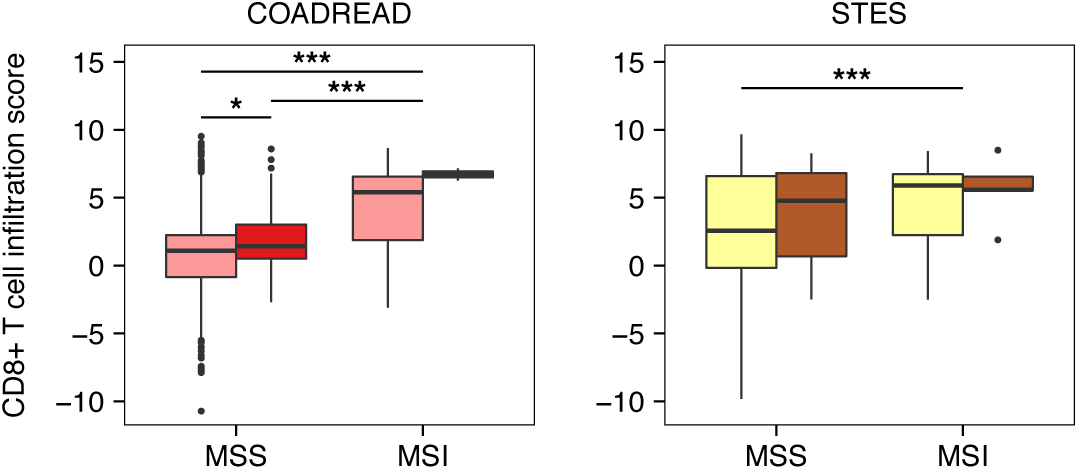
CD8+ T cell infiltration after adjusting for microsatellite instability and virus infection status. Boxplots depicting the distribution of CD8+ T cell infiltration scores across COADREAD and STES samples stratified by MSI status and virus infection status. Dark colors indicate virus-infected samples and light colors indicate non-infected samples. Each box spans quartiles with the lines representing the median CD8+ T cell infiltration score for each group. Whiskers represent absolute range excluding outliers. All outliers were included in the plot. P-values were calculated using the Wilcoxon sum-rank test. Significant associations are marked (* one-sided p < 0.05, *** two-sided p < 0.01).

### Infection of HHV4 is associated with decreased T cell receptor repertoire diversity

We next examined how infection of specific viruses associates with different aspects of the infiltrating TCR repertoire. We hypothesized that the presence of viral proteins in the tumor microenvironment may be associated with a high proportion of viral antigen-specific T cell clones, and thus a less diverse TCR repertoire. To test this hypothesis, we employed the TCR Repertoire Utilities in Solid Tissue (TRUST) method^29^ to call TCR-specific reads from bulk RNAseq reads from each of the six TCGA tumor types in our study. In all tumor types, TCR read abundance was well-correlated with our expression-based measures of CD8+ T cell infiltration (average Spearman ρ = 0.73, range = 0.58-0.82). Additionally, differences in TCR read abundance between infected and non-infected samples was largely consistent with our previous findings, indicating concordance between the computational methods chosen for our infiltration analyses (Supplementary Figure S2). We defined the diversity of each patient’s TCR repertoire as the number of unique clonotypes per thousand mapped TCR reads and compared how this metric differed between infected and non-infected samples across each tumor (Figure 3A). In most cases, there were no significant differences in diversity between subgroups. However, in STES, samples infected with HHV4 exhibited significantly lower levels of TCR diversity, indicating an antigen-driven T cell response (P = 1e-3). Furthermore, in the four other tumor types with at least one HHV4 infection, HHV4-infected samples exhibited the lowest median levels of TCR diversity compared to the other patient subgroups. To provide more power to these analyses, we pooled the associations for each virus across cancer types and calculated and determined significance using a meta-z score approach (Figure 3B). HHV4 remained the only virus associated with differing levels of TCR diversity, with a meta-z-score of −3.81 that corresponded to a significant decrease (two-tailed meta-p-value = 1e-4). These results indicated that the presence of HHV4 viral proteins may elicit a clonal T cell response in different tumor types.

**Figure 3:**
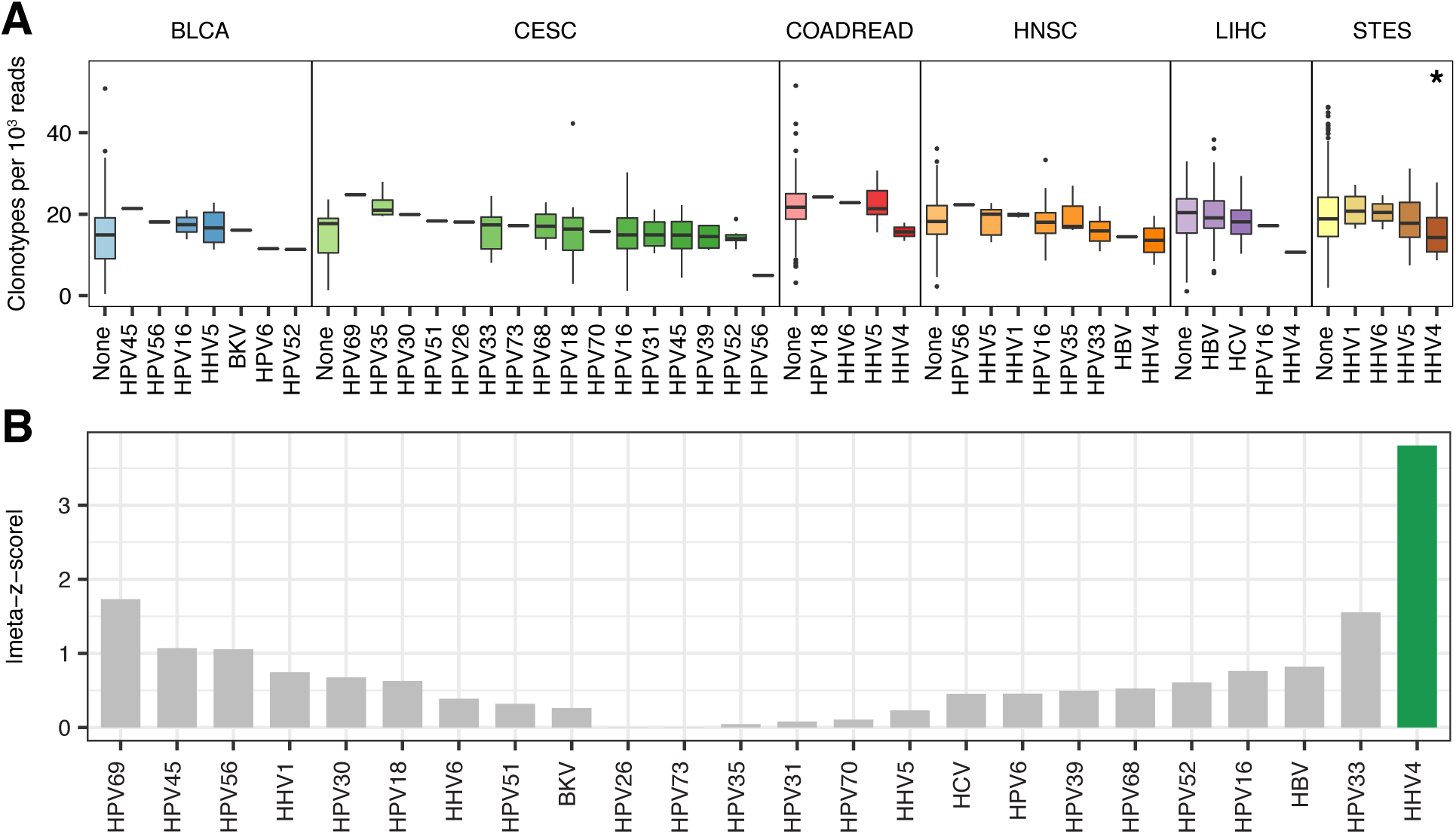
T cell receptor repertoire diversity between samples infected with different virus families. **A,** Boxplots depicting the distribution of unique CDR3 calls (clonotypes) per 1,000 TCR reads in samples from six tumor types infected with different viruses. Each box spans quartiles with the lines representing the median clonal diversity for each group. Whiskers represent absolute range excluding outliers. All outliers were included in the plot. P-values were calculated using the two-tailed Wilcoxon sum-rank test. **B,** Meta-z-score absolute values indicating associations between infection of a given virus and reduced TCR clonal diversity across 6 tumor types. Viruses were ranked by unweighted meta-z-score. Green bars indicate an unweighted meta-z-score < −1.96 (significantly lower TCR clonal diversity, two-tailed p-value = 0.05) while grey bars indicate an unweighted meta-z-score whose absolute value is < 1.96.

To corroborate these findings, we obtained TCR diversity metrics calculated by the MiTCR TCR analysis software from a previous study^32,33^ and compared how each of these metrics, which included Shannon entropy, species richness, and species evenness, varied between samples infected by a given virus and those with no infections (Supplementary Table S2). In agreement with our previous analyses, we found that Shannon entropy and species richness were increased in numerous contexts, such as in HPV-infected CESC and HNSC samples as well as HHV4-infected STES samples, suggesting increased levels of T cell infiltration. Furthermore HHV4-infected STES samples exhibited a moderate reduction in TCR evenness that was indicative of a clonal T cell response (P = 0.08). Together, the MiTCR and TRUST results reinforced each other, providing additional evidence of the altered T cell dynamics associated with virus infection.

### Tissue-specific virus infection gene signatures reproducibly predict infection status

While some gene expression datasets contain immunohistochemistry and sequencing information that can inform a patient’s virus infection status, several do not, making it difficult to further study how virus infection associates with different clinical variables. To address this issue, we devised a method to create a gene expression signature that could predict a patient’s virus infection status. To design this signature, we applied a previously developed approach that weighted each gene in the transcriptome based on how well the gene’s expression level distinguished virus-infected patients from non-infected patients in a generalized linear model.^34^ Genes that more significantly differed between a virus-infected and non-infected sample were given exponentially more weight in the signature than genes that did not significantly distinguish between the two groups of samples. To minimize the effect of confounding on the creation of this signature, each gene’s model was adjusted for age, stage, grade, lymph node metastasis status, and microsatellite instability status depending on the availability of that information.

We first applied this approach to the TCGA HNSC dataset, comparing samples positive for any virus to those negative for all viruses. We then applied the signature back to the TCGA HNSC dataset to calculate a virus infection score for each patient and used these scores to classify whether a sample was virus-positive or virus-negative. The resulting classifications yielded an area under the receiver operating characteristic curve (AUC) of 0.92, indicating high classification accuracy (Figure 4A). To confirm that this high performance was not a result of overfitting in the TCGA dataset, we applied the TCGA-derived signature to a series of additional microarray datasets that also contained clinical measures of virus infection status (Supplementary Table S3). In these datasets, the signature demonstrated excellent accuracy, yielding AUCs ranging from 0.81 to 0.95 (Figure 4B, Supplementary Figure S3). To examine whether this procedure could be used to classify additional tumor types, we derived and tested signatures in in two more cancers, CESC and LIHC, for which there were suitable test datasets for validation. Each tumor type’s respective signature exhibited high accuracy in the dataset from which it was derived (AUC = 0.80 and 0.85 for CESC and LIHC, respectively Figure 4A). In CESC, this performance also translated to an independent dataset (AUC = 0.91). However, in LIHC, the signature performed demonstrably worse in the test dataset (AUC = 0.63), though virus-positive samples still had significantly higher signature scores than virus-negative samples (P = 0.01). These analyses together served as a proof-of-principle that virus status could be inferred from expression information in multiple tumor types.

**Figure 4:**
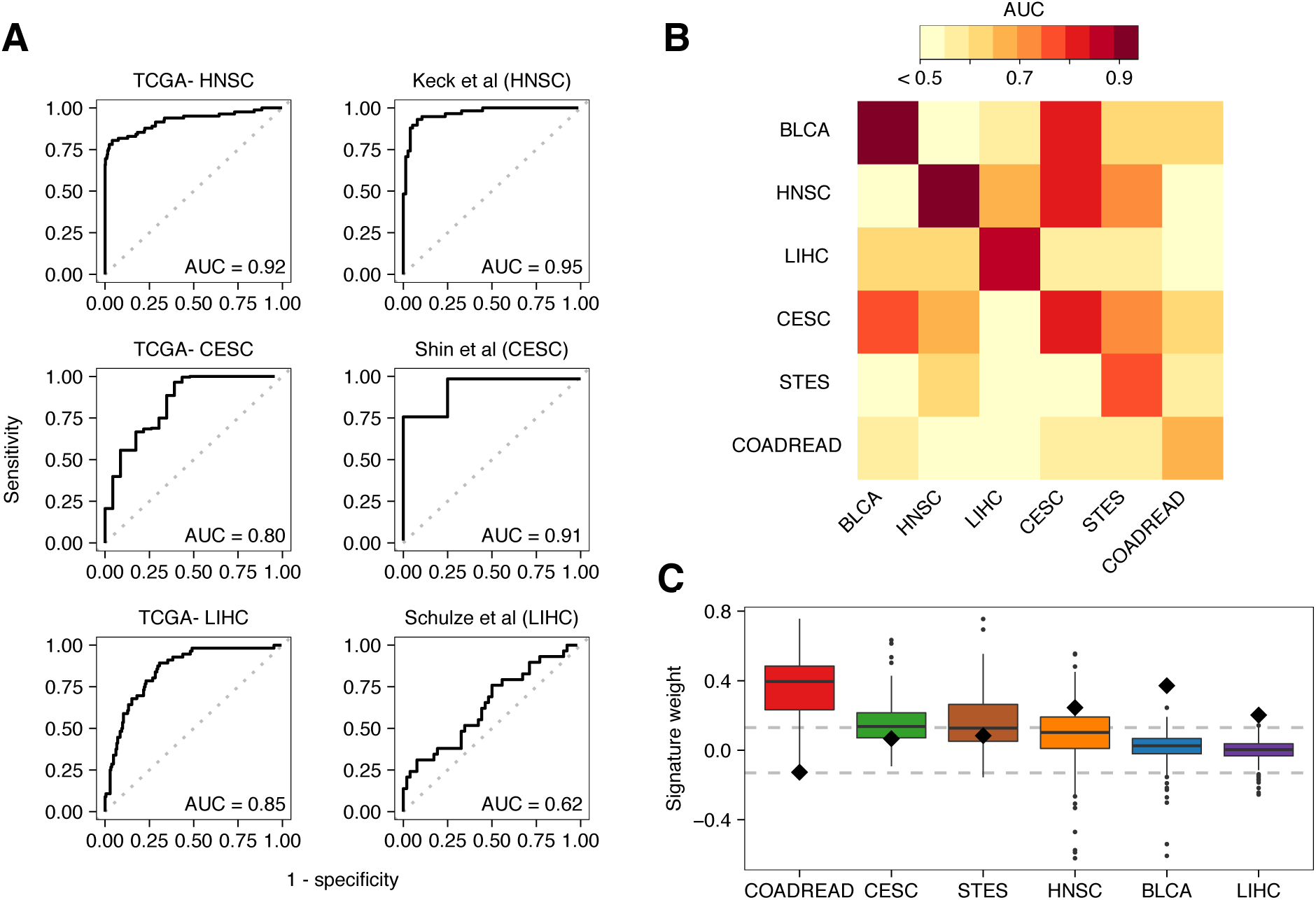
Performance and characterization of the virus infection gene expression signature. **A,** ROC curves illustrating the accuracy of using the virus gene expression signature to classify infected samples from non-infected samples. Plots on the left depict the signature’s performance in training data while plots on the right depict the signature’s performance in test datasets. From top to bottom, test datasets were obtained from GEO under accession numbers GSE40774, GSE49288, and GSE62232. **B,** Heatmap of AUCs for signatures trained in one tissue type (columns) and applied to another (rows). To show contrast, all AUCs < 0.5 were trimmed to 0.5. **C,** Boxplots depicting the weight of the gene *MKI67* (black diamond) and the weight distribution of genes comprising the ESTIMATE immune gene expression signature (boxes). Dotted lines at 0.13 indicate threshold at which weights correspond to P < 0.05. to significant Each box spans quartiles with the lines representing the median signature weight in each group. Whiskers represent absolute range excluding outliers. All outliers were included in the plot.

Given the associations between virus infection and increased immune infiltration across different tumor types, we hypothesized that it might be possible to predict the virus infection using signatures derived from a tumor type different from the cancer to which they were applied. Successful prediction of infection status in a tissue-agnostic manner would suggest a shared biology between tumor types following virus infection. To determine the extent to which this was true, we derived a virus infection signature from each of the six virus-associated TCGA tumor types and then assessed each signature’s performance in cross-tissue classifications using the AUC (Figure 4B). We found unsurprisingly that each signature performed best in the tissue type it was derived from, with AUCs ranging from 0.65 (COADREAD) to 0.94 (BLCA), and a mean of 0.82. However, we also identified four cross-tissue predictions that yielded AUCs greater than the lowest same-tissue AUC, with the BLCA signature performing well in CESC (AUC = 0.78), the CESC signature performing well in BLCA and HNSC (AUC = 0.83 and 0.82, respectively), and the STES signature performing well in HNSC and CESC (AUC = 0.72 and 0.70, respectively).

It has broadly been shown that viruses alter the transcriptional activity of the cells they infect by disrupting cell cycle control and inducing proliferation-associated expression programs.^18^ Furthermore, our previous analyses indicated that the transcriptomes of virus-infected samples are affected by the increased inflammation present in their tumor microenvironments. To better characterize the signals detected by each virus infection signature, we examined how heavily they weighed the single-gene proliferation marker, marker of cell proliferation KI67 (*MKI67*), and the multi-gene ESTIMATE immune signature^35^, which is a well-characterized genomic marker of immune infiltration (Figure 4C). We found that *MKI67* was weighted most highly in BLCA, HNSC, and LIHC, with weights corresponding to significantly higher proliferation rates in virus-infected cancers (P = 2e-4, 3e-3, and 0.01, respectively). Additionally, at least 40% of the ESTIMATE genes in the CESC, COADREAD, HNSC, and STES signatures were weighted at levels that corresponded to significantly higher immune infiltration levels in virus-infected samples (P < 0.05). In BLCA and LIHC, this number was 10% and 4%, respectively. Considering that the virus infection score is exponentially more influenced by significant genes than insignificant ones, these results indicate that the CESC, COADREAD, HNSC, and STES signatures are primarily dominated by immune genes, while the BLCA signature detects a combination of immune and proliferative signals and the LIHC signature is primarily proliferative.

### Expression-inferred virus status is differentially associated with survival across cancers

We questioned how the altered expression programs associated with virus infection affect patient survival. Increased cell proliferation is a well-known negative prognostic factor.^36^ However, immune infiltration has been associated with prolonged survival in a large variety of cancers.^37^ To answer this question, we applied each virus infection gene expression signature to the microarray datasets of the same cancer type in the Prediction of Clinical Outcomes from Genomics (PRECOG) meta-dataset.^38^ We then modeled the relationship between virus score and patient survival in each dataset using a Cox proportional hazards model and pooled the resulting z-scores to get a summary statistic capturing the association between virus infection and patient survival in each tumor type (Supplementary Figure S4). We identified two tumor types, HNSC and BLCA, that exhibited significant meta-associations between tissue-specific virus score and patient survival (meta-p-value < 0.05). In HNSC, this score was reproducibly associated with prolonged patient prognosis. However, in BLCA, this trend was reversed with the BLCA-specific virus score reproducibly associated with shorter survival (Supplementary Table S4). To determine whether these associations were due to the immune- or proliferation-associated programs captured by each tissue’s respective virus signature, we chose one dataset from each tumor type, dichotomized the samples by the median virus score into signature-high and signature-low groups, and then examined how *MKI67* expression and the ESTIMATE immune score differed between the two groups (Figure 5). Notably, these datasets did not include gold standard virus infection information, demonstrating the utility of our signature in expanding genomic studies of virus infection. In HNSC, *MKI67* expression did not differ between signature-high and signature-low patients, while ESTIMATE immune scores were significantly higher in signature-high patients than in signature-low patients (Figure 5B; P = 4e-3). We replicated these associations in an additional HNSC dataset that lacked survival information but had gold-standard virus infection information (Supplementary Figure S5). This suggested that in HNSC, virus infection can induce a prolonged patient survival phenotype by inducing a tumor immune response. In BLCA, signature-high patients exhibited significantly higher levels of *MKI67* expression (P = 3e-4) and ESTIMATE immune signature scores (Figure 5D; P = 1e-5). This finding was interesting, as it indicated that virus infections in BLCA could induce both increased immune infiltration and increased cell proliferation. However, the association between virus infection and shorter survival suggested that the increased proliferation of the tumor cells overwhelms the ability of the immune system to keep tumor development in check.

**Figure 5:**
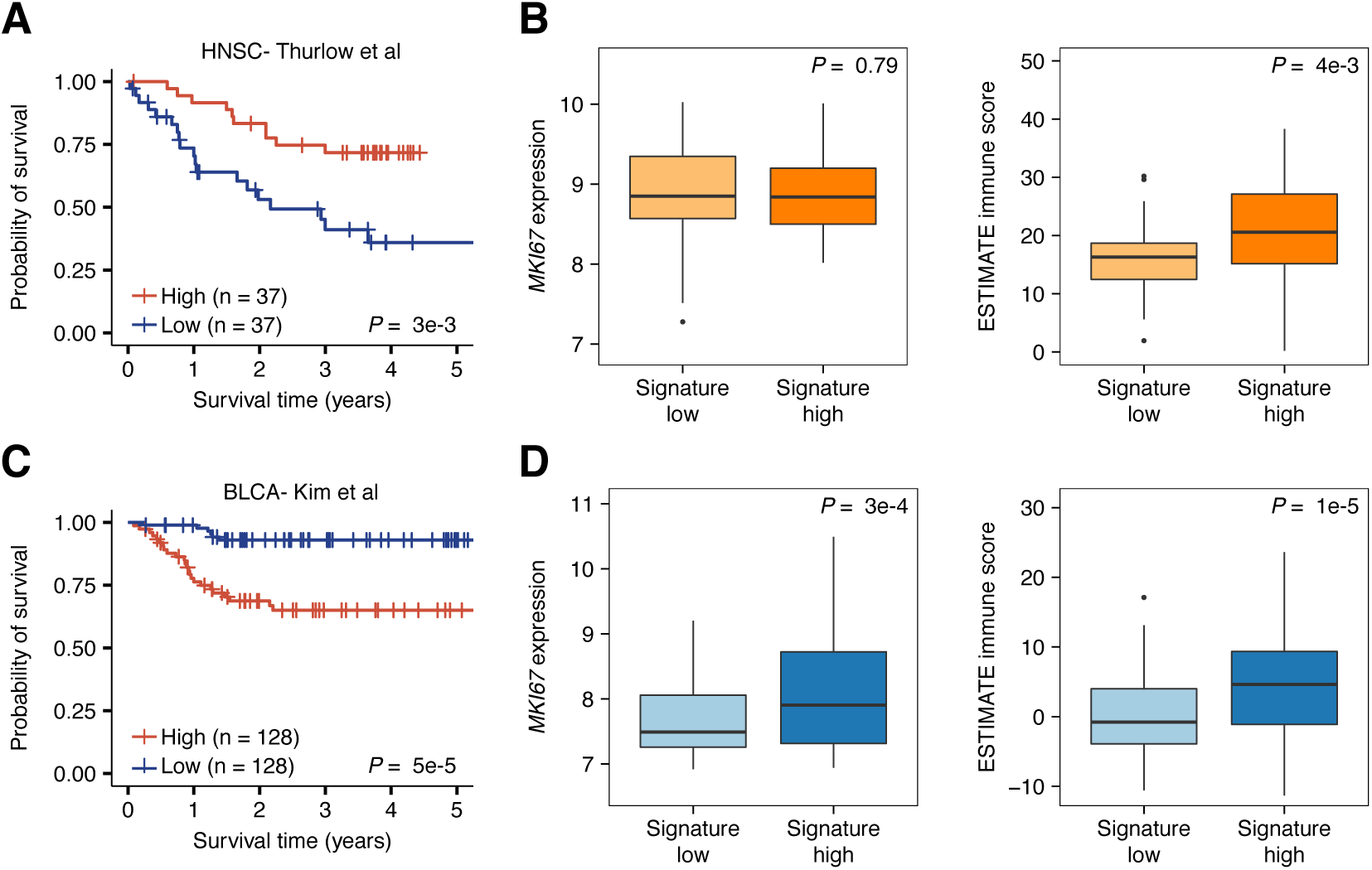
Association between the virus infection gene expression signature and survival of head and neck cancer and bladder patients. **A,** Kaplan-Meier plot depicting the survival probability over five years for samples with high (red) and low (blue) virus infection signature scores in the Thurlow et al head and neck cancer dataset. **B,** Boxplots depicting the difference in *MKI67* expression (left) and ESTIMATE immune score (right) between signature low and signature high samples in the Thurlow et al dataset. **C,** Kaplan-Meier plot depicting the survival probability over time for samples with high (red) and low (blue) virus infection signature scores in the Kim et al bladder cancer dataset. **D,** Boxplots depicting the difference in *MKI67* expression (left) and ESTIMATE immune score (right) between signature low and signature high samples in the Kim et al dataset. For all Kaplan-Meier plots, samples were stratified into high and low groups using the median virus infection score. P-values were calculated using the log-rank test and indicate difference between the survival distributions of the full dataset. Vertical hash marks indicate censored data. In all boxplots, boxes span quartiles with the lines representing the median expression or score for each group. Whiskers represent absolute range excluding outliers. All outliers were included in the plot. P-values were calculated using the two-tailed Wilcoxon sum-rank test.

## Discussion

Virus-induced cancers are distinct from other tumor types in that they are the result of the actions of an infectious agent rather than a mutagenic process. Upon infection, viruses can induce the expression and release of pathogenic proteins into the tumor microenvironment, making the cancers they infect an interesting model for which to investigate the tumor immune response. In this study, we have profiled the tumor microenvironment and genetic programs associated with virus infection in six cancer types. Our analyses provide new insights into how viruses reshape the tumor microenvironment, identifying differences in immune infiltration and TCR diversity in infected compared to non-infected samples. Additionally, our work furthers understanding of how viruses affect tumor proliferation and patient survival by expanding genomic studies of virus infections into additional datasets. Collectively, these results may aid in efforts to identify virus-associated cancer patient subgroups that are sensitive to immunotherapy.

We identified three cancer types, CESC, HNSC, and STES, where the microenvironment of viral protein-expressing samples was consistently more infiltrated than that of their non-infected counterparts. Additionally, when adjusting for microsatellite instability status, we found COADREAD samples showed similar trends. These tumor types are primarily infected by human papilloma viruses and human herpesviruses, which are both known to be involved in direct carcinogenesis. Several studies have characterized the immunogenic nature of these viral proteins, and our results indicate that the presence of these antigens in the tumor microenvironment can elicit a strong immune response.^39-43^ In contrast to these findings, these associations were not present in hepatitis-infected LIHC, despite the fact that the power of our analysis was comparable to that of other cancer types. Hepatitis viruses cause chronic infections of the liver, leading to persistent inflammation that can contribute to the deregulation of the immune response and T cell exhaustion.^44,45^ Our results supported these findings, with hepatitis B virus-infected patients exhibiting decreases in the infiltration of cytolytic cells and the down-regulation of antigen presentation machinery, both hallmarks of an immunosuppressed microenvironment. Together, our results illustrate how differences in viral prevalence across different tissue types may shape our understanding of the tumor microenvironment of the cancer at the population level. Virus infection should thus be carefully considered when studying the tumor-immune interactions of these cancers.

To better understand how viral proteins can alter the adaptive immune response in these cancers, we examined how the clonal dynamics of the infiltrating TCR repertoire varied across cancers infected with different viruses. Interestingly, HHV4-infected samples exhibited the lowest TCR diversity levels in STES and other tumor types, suggesting evidence of a clonal T cell response. Intriguingly, a recent study has identified a gastric cancer patient positive for HHV4 that exhibited clinical benefit in response to the anti-PD-L1 drug avelumab.^23^ Furthermore, this study found, in agreement with our findings, that HHV4-positive patients exhibit increased immune infiltration. Given that high immune infiltration and low TCR diversity have both been associated with response to immune checkpoint blockade^14,46^, our analyses suggest that HHV4-positive cancers may represent good targets for future immunotherapeutic approaches. Beyond HHV4, we observed no significant differences in TCR diversity between samples infected with a given virus and non-infected samples. While surprising, this result is in agreement with a study in CESC, which found that the primary antitumor T cell responses in patients receiving adoptive T cell therapy are directed against tumor-intrinsic neoantigens or cancer germline antigens, rather than viral antigens.^24^ Going forward, analyses performing targeted TCR sequencing on patient samples may further our understanding of how virus infection alters the clonal dynamics of the T cell response. This will be important, as our early results suggest that improving our understanding of this process can further refine our ability to identify immunotherapy-sensitive patients.

To complement our studies of the virus-associated tumor microenvironment, we developed a series of tissue-specific virus infection signatures capable of predicting a patient’s virus infection status from their gene expression profile. These signatures were highly accurate in a variety of contexts, with virus-infected patients exhibiting significantly higher scores than non-infected ones. Functional characterization of these signatures revealed that the CESC, COADREAD, HNSC, and STES were associated with immune-based signals, while LIHC and BLCA were more indicative of higher cell proliferation rates. These results suggested that viruses of different families can induce distinct genetic reprogramming in the tumors they infect. Survival analyses using these signatures supported this finding, with virus-high HNSC patients exhibiting a prolonged prognosis and high levels of immune infiltration while virus-high BLCA patients exhibiting shorter survival and higher cell proliferation rates, despite also having a higher immune infiltration level. The similarities and differences in the signatures across cancer types are likely due to the viruses that are most prevalent in them, and our results thus indicate the importance of understanding how infections by different viruses can affect tumor development.

In summary, we present a multi-tissue analysis of the microenvironment and genetic changes associated with virus infection. Our results highlight the divergent changes associated with virus infection in different tumor types and suggest that viruses of different families should be treated as unique features with regards to the design of immunotherapeutic approaches. Going forward, it will be important to collect genomic data and virus infection information from additional patients., as in several cancer types infection is exceedingly rare and power to detect significant associations is limited. By integrating this information with findings from previous studies, we can begin to fully appreciate the role of viruses in driving tumor progression and harness this knowledge for therapeutic benefit.

## Methods

### Datasets

Virus abundance information, quantified as the number of virus supporting reads per hundred million reads processed (RPHM) for 2,343 virus-associated TCGA tumor was downloaded from a supplementary file from a previous study.^26^ Of the samples included in this dataset, 2,341 had matching RNAseq gene expression information, which was obtained from GDAC FireHose (RNAseqV2, RSEM). TCGA RNAseqV2 data was log10-transformed for infiltration score calculation. For patients with multiple samples, the average RSEM value for each gene was used. Raw RNAseq paired-end reads (.fastq) for TCGA samples were downloaded from the Genomic Data Commons legacy archive (https://portal.gdc.cancer.gov/legacy-archive). RNAseq reads were aligned to human reference genome hg19 using Bowtie2^47^ run with default parameters. MANTIS microsatellite instability scores for TCGA samples were downloaded from a previous publication.^48^ All TCGA sample size information is available as a supplement (Supplementary Table S1). Additional gene expression data and the associated virus infection and survival information were obtained from the gene expression omnibus (GEO) under accession numbers GSE40774, GSE6791, GSE55550, GSE39366, GSE65858, GSE49288, GSE62232, GSE44001, as well as from PRECOG (https://precog.stanford.edu/precog_data.php; Supplementary Table S3).^38^

Based on findings from the original study, all TCGA samples exhibiting ≥ 100 RPHM for a given virus were classified as infected by that virus. In the event that a sample was positive for multiple viruses, we classified the sample as positive for the virus that exhibited the highest level of RPHM. For microsatellite instability, binary thresholds were determined from the distribution of MANTIS scores in each cancer type, with the default of 0.4 used for STES and 0.5 used for COADREAD.

### Calculation of immune infiltration scores and T cell receptor profiling

Immune infiltration scores for CD8+ T cells, B cells, NK cells, and macrophages were calculated as described previously using the same four previously validated signatures derived from the Immunological Genome Project.^27^ T cell receptor profiling was run on Bowtie2-aligned TCGA RNAseq reads (.bam) using TRUST version 3.0 (https://bitbucket.org/liulab/trust/).^29^ During the alignment step, TRUST requires for unmapped reads to be included, local alignment to be disabled, and for the number of mismatches tolerated from mapped reads to be no greater than 2, all default parameters of Bowtie2. TCR clonotype diversity was estimated using the clonotypes per thousand mapped TCR reads for each sample. This number was calculated from TRUST’s output by dividing each sample’s number of unique CDR3 reads (denoted as “cdr3dna” in TRUST’s tabular output) by its total number of TCR reads (denoted as “est_lib_size”) divided by 1,000. ESTIMATE immune scores were calculated using the ESTIMATE R package (http://bioinformatics.mdanderson.org/estimate/index.html).

### Derivation and application of the virus infection gene expression signature

The virus infection gene expression signature was designed to capture the transcriptome-wide differential gene expression activity between virus-positive and virus-negative patients within a given TCGA tumor type. To define this signature, a logistic regression model was constructed for each gene in the transcriptome with a patient’s virus infection status as the response variable and the expression level of that gene as the predictor variable. To ensure the signature most accurately captured the difference between virus-positive and virus-negative samples, potential confounding factors, including stage, age, grade, lymph node metastasis status, and microsatellite instability status were also included in the model as covariates. A covariate was only included in a specific cancer type’s model when less than half of the samples of that cancer type had NA values for that covariate, and in the case of categorical variables, when fewer than 80% of samples were the same category for that variable (Supplementary Table S5). The model used for each gene can be formulized below, where Y is a patient’s virus infection status (1 indicating positive, 0 indicating negative), X_1_ is the expression of the gene under consideration and X_2_ through X_n_ are the n-1 covariates:

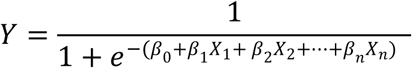

From these logistic regression models, we derived each gene’s coefficient (β-value), which indicated whether the gene was up- or down-regulated in virus-positive samples compared to virus-negative samples, as well as the p-value of the association between each gene’s expression level and virus infection status. We then used these statistics to define the final virus infection gene expression signature as a set of gene-specific weight profiles that indicated the magnitude and directionality (up-regulated or down-regulated) of the association between a gene’s expression level and a patient’s virus infection status. In the up-regulated weight profile, the p-values of all genes with a β-coefficient > 0 were -log10-transformed and the remainder were set to 0, while in the down-regulated weight profile, the p-values of all genes with a β-coefficient < 0 were -log10-transformed while the remaining p-values were set to 0. The resulting numbers > 10 were trimmed to 10 to avoid outliers and then all numbers were rescaled to between 0 and 1 to obtain the final weight signatures.

To calculate the virus infection scores in a series of patients, these signatures along with a patient gene expression dataset were input into the Binding Association with Sorted Expression (BASE) algorithm.^49^ Briefly, BASE functions by examining how the weights of each input signature are distributed through a patient’s ranked gene expression profile. Patients that exhibit high expression of genes with high up-regulated weights and low expression of genes with high down-regulated weights are assigned a higher score for that signature, while patients deviating from this pattern are assigned a lower score. Full details on how BASE uses these signatures to calculate the score are available in a previous study.^34^ The virus signatures formatted for BASE input are available as a supplement (Supplementary Table S5).

To evaluate the accuracy of the signature, we ranked each patient by their virus infection scores and then performed an iterative procedure where each patient’s score in the ranked list was used as a threshold by which to classify patients as virus-positive or virus-negative. For each iteration, we calculated the sensitivity and specificity of the resulting classification and then used these numbers to calculate the AUC.

### Statistical analyses

All comparisons of the distributions between two groups were made using the two-tailed Wilcoxon-sum rank test unless stated otherwise. Logistic regression modeling to derive the virus signature was performed using the “glm()” function in R with family “binomial.” For two-class survival comparisons, samples were stratified into high and low groups based on whether they were above or below the median virus score. Statistical significance of the difference between these survival distributions was calculated using the log-rank test through the “survdiff()” function from the R survival package. Cox proportional hazards regression was used to model the association between virus score as a continuous variable and patient survival, and was performed using the “coxph()” function from the R survival package. For meta-analyses, p-values derived from Wilcoxon sum-rank tests or Cox proportional hazards models were converted to z-scores. The z-scores were then collapsed into meta-z-scores by applying an unweighted version of Stouffer’s method.^50^

## Acknowledgments

We thank Randolph Noelle for his valuable suggestions in editing this manuscript. This work was supported by the National Institutes of Health under award number 1R01CA547195, the National Center for Advancing Translational Sciences of the National Institutes of Health under award number UL1TR001086, the Norris Cotton Cancer Center Core Grant Developmental Funds, and the Geisel School of Medicine at Dartmouth College start-up funding package provided to C.C.

## Author Contributions

C.C. directed the project. F.S.V. and C.C. conceived the research and designed the method and experiments. F.S.V., Y.W., and C.C. curated the data. F.S.V. performed the analysis and produced the figures. F.S.V., Y.W., and C.C. interpreted the results. F.S.V. drafted the manuscript. F.S.V., Y.W., and C.C. read and approved the final manuscript.

## Competing Financial Interests

The authors declare no competing financial interests.

## References

1 Pardoll, D. M. The blockade of immune checkpoints in cancer immunotherapy. Nat Rev Cancer 12, 252-264, doi:10.1038/nrc3239 (2012).

2 Alsaab, H. O. et al. PD-1 and PD-L1 Checkpoint Signaling Inhibition for Cancer Immunotherapy: Mechanism, Combinations, and Clinical Outcome. Front Pharmacol 8, 561, doi:10.3389/fphar.2017.00561 (2017).

3 Robert, C. et al. Nivolumab in previously untreated melanoma without BRAF mutation. N Engl J Med 372, 320-330, doi:10.1056/NEJMoa1412082 (2015).

4 Borghaei, H. et al. Nivolumab versus Docetaxel in Advanced Nonsquamous Non-Small-Cell Lung Cancer. N Engl J Med 373, 1627-1639, doi:10.1056/NEJMoa1507643 (2015).

5 Garon, E. B. et al. Pembrolizumab for the treatment of non-small-cell lung cancer. N Engl J Med 372, 2018-2028, doi:10.1056/NEJMoa1501824 (2015).

6 Motzer, R. J. et al. Nivolumab versus Everolimus in Advanced Renal-Cell Carcinoma. N Engl J Med 373, 1803-1813, doi:10.1056/NEJMoa1510665 (2015).

7 Hodi, F. S. et al. Improved survival with ipilimumab in patients with metastatic melanoma. N Engl J Med 363, 711-723, doi:10.1056/NEJMoa1003466 (2010).

8 Schumacher, T. N. & Schreiber, R. D. Neoantigens in cancer immunotherapy. Science 348, 69-74, doi:10.1126/science.aaa4971 (2015).

9 Turajlic, S. et al. Insertion-and-deletion-derived tumour-specific neoantigens and the immunogenic phenotype: a pan-cancer analysis. Lancet Oncol 18, 1009-1021, doi:10.1016/S1470-2045(17)30516-8 (2017).

10 Goodman, A. M. et al. Tumor Mutational Burden as an Independent Predictor of Response to Immunotherapy in Diverse Cancers. Mol Cancer Ther 16, 2598-2608, doi:10.1158/1535-7163.MCT-17-0386 (2017).

11 Snyder, A. et al. Genetic basis for clinical response to CTLA-4 blockade in melanoma. N Engl J Med 371, 2189-2199, doi:10.1056/NEJMoa1406498 (2014).

12 Rizvi, N. A. et al. Cancer immunology. Mutational landscape determines sensitivity to PD-1 blockade in non-small cell lung cancer. Science 348, 124-128, doi:10.1126/science.aaa1348 (2015).

13 Le, D. T. et al. PD-1 Blockade in Tumors with Mismatch-Repair Deficiency. N Engl J Med 372, 2509-2520, doi:10.1056/NEJMoa1500596 (2015).

14 Van Allen, E. M. et al. Genomic correlates of response to CTLA-4 blockade in metastatic melanoma. Science 350, 207-211, doi:10.1126/science.aad0095 (2015).

15 Rosenberg, J. E. et al. Atezolizumab in patients with locally advanced and metastatic urothelial carcinoma who have progressed following treatment with platinum-based chemotherapy: a single-arm, multicentre, phase 2 trial. Lancet 387, 1909-1920, doi:10.1016/S0140-6736(16)00561-4 (2016).

16 Johnson, D. B. et al. Targeted Next Generation Sequencing Identifies Markers of Response to PD-1 Blockade. Cancer Immunol Res 4, 959-967, doi:10.1158/2326-6066.CIR-16-0143 (2016).

17 Vandeven, N. & Nghiem, P. Pathogen-driven cancers and emerging immune therapeutic strategies. Cancer Immunol Res 2, 9-14, doi:10.1158/2326-6066.CIR-13-0179 (2014).

18 Moore, P. S. & Chang, Y. Why do viruses cause cancer? Highlights of the first century of human tumour virology. Nat Rev Cancer 10, 878-889, doi:10.1038/nrc2961 (2010).

19 Rooney, M. S., Shukla, S. A., Wu, C. J., Getz, G. & Hacohen, N. Molecular and genetic properties of tumors associated with local immune cytolytic activity. Cell 160, 48-61, doi:10.1016/j.cell.2014.12.033 (2015).

20 Li, B. et al. Comprehensive analyses of tumor immunity: implications for cancer immunotherapy. Genome Biol 17, 174, doi:10.1186/s13059-016-1028-7 (2016).

21 Seiwert, T. Y. et al. Safety and clinical activity of pembrolizumab for treatment of recurrent or metastatic squamous cell carcinoma of the head and neck (KEYNOTE-012): an open-label, multicentre, phase 1b trial. Lancet Oncol 17, 956-965, doi:10.1016/S1470-2045(16)30066-3 (2016).

22 Nghiem, P. T. et al. PD-1 Blockade with Pembrolizumab in Advanced Merkel-Cell Carcinoma. N Engl J Med 374, 2542-2552, doi:10.1056/NEJMoa1603702 (2016).

23 Panda, A. et al. Immune Activation and Benefit From Avelumab in EBV-Positive Gastric Cancer. J Natl Cancer Inst 110, 316-320, doi:10.1093/jnci/djx213 (2018).

24 Stevanovic, S. et al. Landscape of immunogenic tumor antigens in successful immunotherapy of virally induced epithelial cancer. Science 356, 200-205, doi:10.1126/science.aak9510 (2017).

25 Tang, K. W., Alaei-Mahabadi, B., Samuelsson, T., Lindh, M. & Larsson, E. The landscape of viral expression and host gene fusion and adaptation in human cancer. Nat Commun 4, 2513, doi:10.1038/ncomms3513 (2013).

26 Cao, S. et al. Divergent viral presentation among human tumors and adjacent normal tissues. Sci Rep 6, 28294, doi:10.1038/srep28294 (2016).

27 Varn, F. S., Wang, Y., Mullins, D. W., Fiering, S. & Cheng, C. Systematic Pan-Cancer Analysis Reveals Immune Cell Interactions in the Tumor Microenvironment. Cancer Res 77, 1271-1282, doi:10.1158/0008-5472.CAN-16-2490 (2017).

28 Newman, A. M. et al. Robust enumeration of cell subsets from tissue expression profiles. Nat Methods 12, 453-457, doi:10.1038/nmeth.3337 (2015).

29 Li, B. et al. Landscape of tumor-infiltrating T cell repertoire of human cancers. Nat Genet 48, 725-732, doi:10.1038/ng.3581 (2016).

30 Mose, L. E. et al. Assembly-based inference of B-cell receptor repertoires from short read RNA sequencing data with V’DJer. Bioinformatics 32, 3729-3734, doi:10.1093/bioinformatics/btw526 (2016).

31 Hause, R. J., Pritchard, C. C., Shendure, J. & Salipante, S. J. Classification and characterization of microsatellite instability across 18 cancer types. Nat Med 22, 1342-1350, doi:10.1038/nm.4191 (2016).

32 Bolotin, D. A. et al. MiTCR: software for T-cell receptor sequencing data analysis. Nat Methods 10, 813-814, doi:10.1038/nmeth.2555 (2013).

33 Thorsson, V. et al. The Immune Landscape of Cancer. Immunity 48, 812-830 e814, doi:10.1016/j.immuni.2018.03.023 (2018).

34 Zhao, Y. et al. A P53-Deficiency Gene Signature Predicts Recurrence Risk of Patients with Early-Stage Lung Adenocarcinoma. Cancer Epidemiol Biomarkers Prev 27, 86-95, doi:10.1158/1055-9965.EPI-17-0478 (2018).

35 Yoshihara, K. et al. Inferring tumour purity and stromal and immune cell admixture from expression data. Nat Commun 4, 2612, doi:10.1038/ncomms3612 (2013).

36 Hanahan, D. & Weinberg, R. A. Hallmarks of cancer: the next generation. Cell 144, 646-674, doi:10.1016/j.cell.2011.02.013 (2011).

37 Fridman, W. H., Pages, F., Sautes-Fridman, C. & Galon, J. The immune contexture in human tumours: impact on clinical outcome. Nat Rev Cancer 12, 298-306, doi:10.1038/nrc3245 (2012).

38 Gentles, A. J. et al. The prognostic landscape of genes and infiltrating immune cells across human cancers. Nat Med 21, 938-945, doi:10.1038/nm.3909 (2015).

39 Chen, L. P., Thomas, E. K., Hu, S. L., Hellstrom, I. & Hellstrom, K. E. Human papillomavirus type 16 nucleoprotein E7 is a tumor rejection antigen. Proc Natl Acad Sci U S A 88, 110-114 (1991).

40 Ramos, C. A. et al. Human papillomavirus type 16 E6/E7-specific cytotoxic T lymphocytes for adoptive immunotherapy of HPV-associated malignancies. J Immunother 36, 66-76, doi:10.1097/CJI.0b013e318279652e (2013).

41 Steele, J. C. et al. T-cell responses to human papillomavirus type 16 among women with different grades of cervical neoplasia. Br J Cancer 93, 248-259, doi:10.1038/sj.bjc.6602679 (2005).

42 Merlo, A. et al. The interplay between Epstein-Barr virus and the immune system: a rationale for adoptive cell therapy of EBV-related disorders. Haematologica 95, 1769-1777, doi:10.3324/haematol.2010.023689 (2010).

43 La Rosa, C. & Diamond, D. J. The immune response to human CMV. Future Virol 7, 279-293, doi:10.2217/fvl.12.8 (2012).

44 Seeger, C. & Mason, W. S. Hepatitis B virus biology. Microbiol Mol Biol Rev 64, 51-68 (2000).

45 Ye, B. et al. T-cell exhaustion in chronic hepatitis B infection: current knowledge and clinical significance. Cell Death Dis 6, e1694, doi:10.1038/cddis.2015.42 (2015).

46 Tumeh, P. C. et al. PD-1 blockade induces responses by inhibiting adaptive immune resistance. Nature 515, 568-571, doi:10.1038/nature13954 (2014).

47 Langmead, B. & Salzberg, S. L. Fast gapped-read alignment with Bowtie 2. Nat Methods 9, 357-359, doi:10.1038/nmeth.1923 (2012).

48 Bonneville, R. et al. Landscape of Microsatellite Instability Across 39 Cancer Types. JCO Precision Oncology, 1-15, doi:10.1200/po.17.00073 (2017).

49 Cheng, C., Yan, X., Sun, F. & Li, L. M. Inferring activity changes of transcription factors by binding association with sorted expression profiles. BMC Bioinformatics 8, 452, doi:10.1186/1471-2105-8-452 (2007).

50 Samuel A. Stouffer, E. A. S., Leland C. Devinney, Robin M. Williams Jr. The American Soldier: Adjustment During Army Life. (Princeton University Press, 1949).

