## Supplementary Materials for "Genomic characterization of six virus-associated cancers identifies changes in the tumor microenvironment and altered genetic programming" (from another version of this preprint: version 1, 2018-03-27)

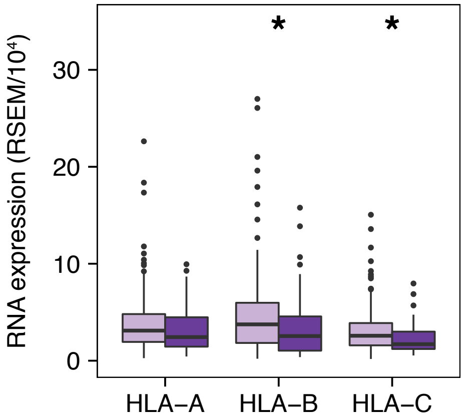


**Supplementary Figure S1: Differences in the expression of HLA genes between HBV-positive and HBV-negative LIHC patients.** Boxplots depicting the distribution of RNA expression for the HLA-A, HLA-B, and HLA-C genes in LIHC samples stratified by HBV-infection status. Y-axis indicates the RSEM-derived RNA expression values divided by 10,000. Dark colors indicate HBV-positive samples and light colors indicate HBV-negative samples. Each box spans quartiles with the lines representing the median clonal diversity for each group. Whiskers represent absolute range excluding outliers. All outliers were included in the plot. Significant associations are marked (* $p < 0.05$, Wilcoxon sum-rank test).

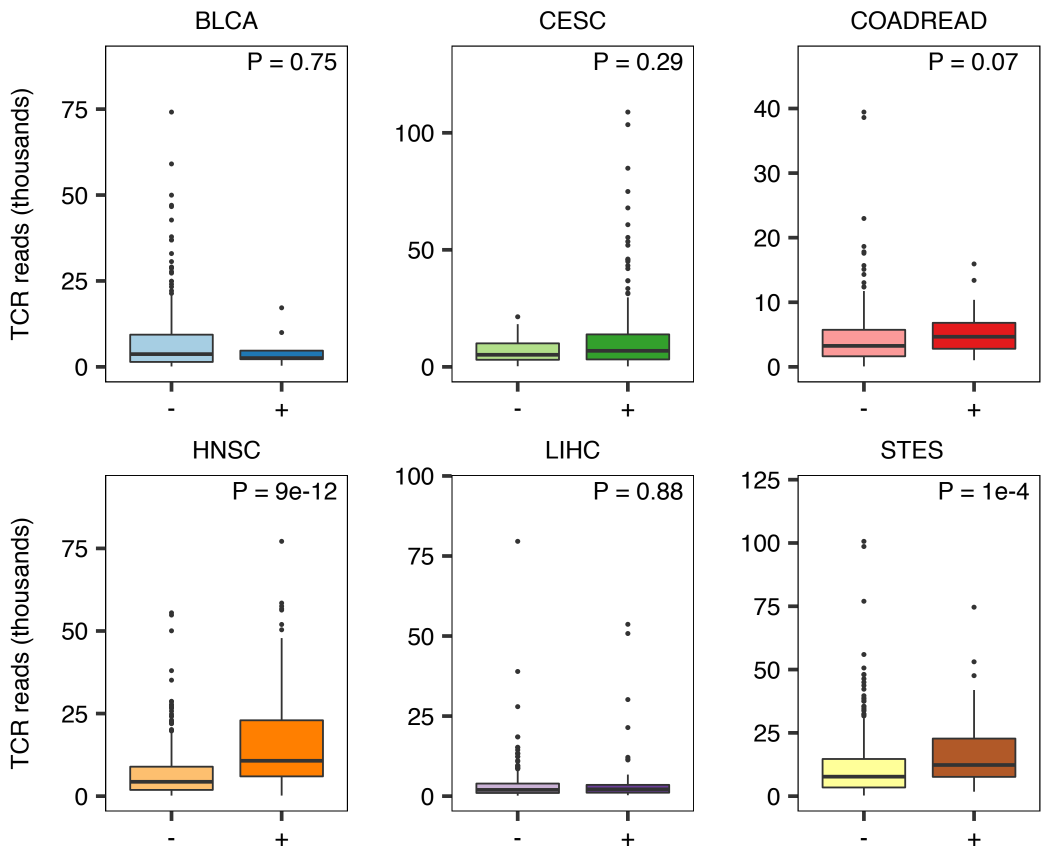


**Supplementary Figure S2: Differences in TCR read abundance between virus-infected and non-infected cancer patients.** Boxplots depicting the distribution of TRUST-inferred TCR reads across samples from six cancer types stratified by virus infection status. Y-axis indicates the number of TCR reads divided by 1,000. Dark colors indicate virus-infected samples and light colors indicate non-infected samples. Each box spans quartiles with the lines representing the median clonal diversity for each group. Whiskers represent absolute range excluding outliers. All outliers were included in the plot. P-values were calculated using the Wilcoxon sum-rank test.

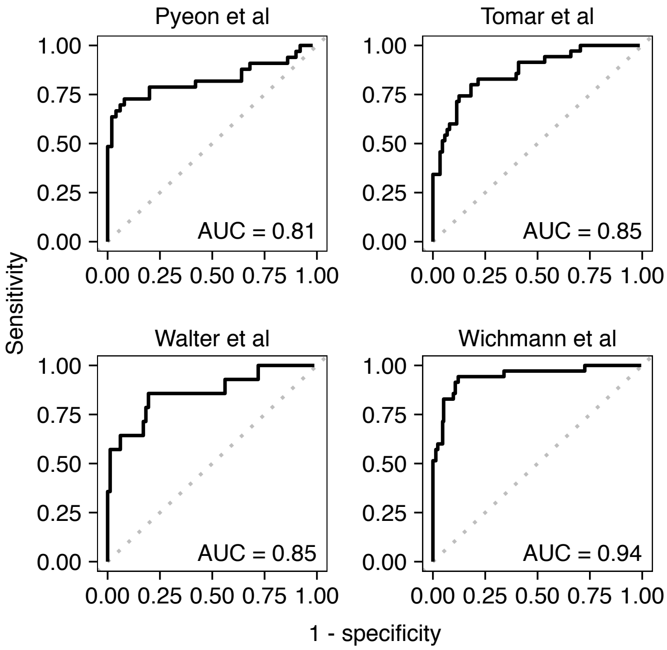


**Supplementary Figure S3: Performance of the virus infection gene expression signature in four HNSC datasets.** ROC curves illustrating the accuracy of using the HNSC-derived virus gene expression signature to classify infected samples from non-infected HNSC samples. The GEO accession numbers for the plots from left to right are GSE6791, GSE55550, GSE39366, and GSE65858.

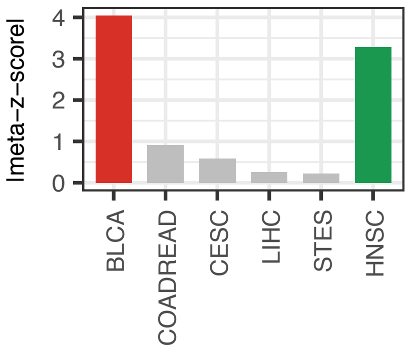


**Supplementary Figure S4: Survival meta-analysis of the virus infection gene expression signature in six cancer types.** Meta-z-score absolute values indicating associations between tissue-specific virus infection score and patient survival across 6 tumor types. Cancers were ranked by unweighted meta-z-score. Red bars indicate cancers with an unweighted meta-z-score > 1.96 (significantly shorter prognosis, two-tailed p-valued = 0.05), grey bars indicate cancers an unweighted meta-z-score whose absolute value is < 1.96 (two-tailed p-value > 0.05), and green bars indicate cancers with an unweighted meta-z-score < -1.96 (significantly prolonged prognosis, two-tailed p-value = 0.05). Datasets used in this meta-analysis were obtained from PRECOG (https://precog.stanford.edu/precog_data.php), with an additional CESC dataset obtained from GEO under accession number GSE44001.

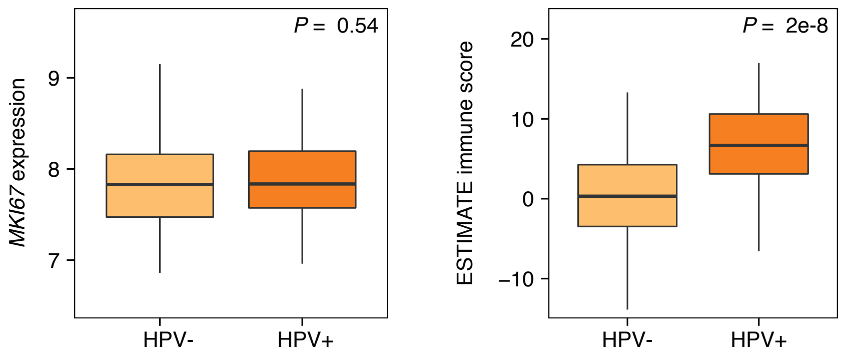


**Supplementary Figure S5: Differences in *MKI67* gene expression and ESTIMATE immune scores between HPV-positive and HPV-negative HNSC patients.** Boxplots depicting the difference in *MKI67* expression (left) and ESTIMATE immune score (right) between HPV-negative and HPV-positive samples in the Keck et al dataset (GSE40774). In all boxplots, boxes span quartiles with the lines representing the median expression or score for each group. Whiskers represent absolute range excluding outliers. All outliers were included in the plot. P-value was calculated using the Wilcoxon sum-rank test.
